# Ancient Genomics Reveals Four Prehistoric Migration Waves into Southeast Asia

**DOI:** 10.1101/278374

**Authors:** Hugh McColl, Fernando Racimo, Lasse Vinner, Fabrice Demeter, J. Víctor Moreno Mayar, Uffe Gram Wilken, Andaine Seguin-Orlando, Constanza de la Fuente Castro, Sally Wasef, Ana Prohaska, Ashot Margarayan, Peter de Barros Damgaard, Rasmi Shoocongdej, Viengkeo Souksavatdy, Thongsa Sayavongkhamdy, Mohd Mokhtar Saidin, Supannee Kaewsutthi, Patcharee Lertrit, Huong Mai Nguyen, Hsiao-chun Hung, Thi Minh Tran, Huu Nghia Truong, Shaiful Shahidan, Ketut Wiradnyana, Anne-Marie Bacon, Philippe Duringer, Jean-Luc Ponche, Laura Shackelford, Elise Patole-Edoumba, Anh Tuan Nguyen, Bérénice Bellina-Pryce, Jean-Christophe Galipaud, Rebecca Kinaston, Hallie Buckley, Christophe Pottier, Simon Rasmussen, Tom Higham, Robert A. Foley, Marta Mirazón Lahr, Ludovic Orlando, Martin Sikora, Charles Higham, David M. Lambert, Eske Willerslev

## Abstract

Two distinct population models have been put forward to explain present-day human diversity in Southeast Asia. The first model proposes long-term continuity (Regional Continuity model) while the other suggests two waves of dispersal (Two Layer model). Here, we use whole-genome capture in combination with shotgun sequencing to generate 25 ancient human genome sequences from mainland and island Southeast Asia, and directly test the two competing hypotheses. We find that early genomes from Hoabinhian hunter-gatherer contexts in Laos and Malaysia have genetic affinities with the Onge hunter-gatherers from the Andaman Islands, while Southeast Asian Neolithic farmers have a distinct East Asian genomic ancestry related to present-day Austroasiatic-speaking populations. We also identify two further migratory events, consistent with the expansion of speakers of Austronesian languages into Island Southeast Asia ca. 4 kya, and the expansion by East Asians into northern Vietnam ca. 2 kya. These findings support the Two Layer model for the early peopling of Southeast Asia and highlight the complexities of dispersal patterns from East Asia.

## Main Text

The population history of Southeast Asia (SEA) has been shaped by interchanging periods of isolation and connectivity. Anatomically modern humans first colonized SEA at least 70,000 years ago (kya) (*1*–*3*). Within SEA, the complex topography and changes in sea level promoted regional expansions and contractions of populations. By the late Pleistocene/early Holocene, a pan-regional lithic technological culture was established across mainland SEA, named Hoabinhian (*4*–*7*). Hoabinhian foragers are thought to be the ancestors of present-day SEA hunter-gatherers, sometimes referred to as ‘Negritos’ because of their comparatively darker skin colour and short stature. Today, however, the majority of people in SEA are believed to be descendants of rice and millet farmers with varying degrees of East Asian phenotypic affinity, suggesting that human diversity in SEA was strongly influenced by population expansions from the north (*4*). Yet, the extent to which the movements from East Asia (EA) impacted on the genetic and cultural makeup of the people of SEA remains controversial.

Two distinct population models have been proposed to account for the biological and cultural diversity of human populations in present-day SEA. The Regional Continuity model, based primarily on morphological evidence, argues for a long-standing evolutionary continuity without significant external gene flow and for the Neolithic transition in SEA occurring as hunter-gatherer groups adopted agriculture, either independently or through cultural contact (*8*–*21*). While this model does not necessarily argue for the independent domestication of crops across SEA, it posits that gene flow from EA farmers was not the main mechanism behind the Neolithic transition. In contrast, the Two Layer model advocates for two major dispersal waves into SEA, where EA farmers replaced the original Hoabinhian inhabitants across SEA through a major demographic southward expansion *ca.* 4 kya. The exception to this would be the isolated populations of the Andaman Islands, peninsular Thailand/Malaysia and the Philippines which are considered the primary descendants of Hoabinhian hunter-gatherers (*22, 23*). Under this model, the migratory wave of farmers originated in present-day China, where rice and millet were fully domesticated in the Yangtze and Yellow River valleys between 9-5.5 kya, and paddy fields developed by 4.5 kya (*4, 24*–*26*). Farming practices are thought to have accompanied these populations as they spread southward through two main routes – an inland wave associated with the expansion of Austroasiatic languages, and an island-hopping route associated with Austronesian languages which eventually reached the Pacific (*27, 28*). Within mainland SEA (MSEA), exchanges with EA appear to have continued in the recent past, however, the extent to which these expansions had a genetic impact on the indigenous populations is unknown.

Genetic studies of contemporary SEA populations have not resolved these controversies (*29*–*32*). Ancient genomics can provide direct evidence of past population events. However, SEA is characterised by tropical and monsoonal climates which cause heavy weathering and acidification of soils (*33*), so ancient genomic studies have, so far, been unsuccessful there. Though shotgun sequencing has revolutionized ancient genomic studies by allowing the retrieval of all mappable DNA fragments from an ancient sample (*34, 35*), the inverse relationship between the proportion of endogenous DNA and the cost of shotgun sequencing makes this approach impractical to apply widely to regions with poor DNA preservation such as SEA. Genome wide SNP capture is one way to circumvent the issue (*36, 37*), but it retrieves only a pre-selected subset of all variants of the genome and thus sacrifices the full potential of rare and irreplaceable fossil samples. An alternative approach is whole genome capture in which human ancient human DNA fragments are enriched through hybridisation to baits that cover the entire mappable human genome (*15*).

We performed comparative testing of three different capture approaches for human DNA - the SeqCap EZ Human Exome Kit v3.0 cat no. 6740294001 (Roche Nimblegen, CA, USA), the SureSelect Human All Exon V5+UTRs cat. no. 5190-6213 (Agilent Technologies) and the Custom MYbaits Whole Genome Enrichment (WGE) Kit version 2.0 (Arbor Biosciences) - with the aim of applying the most effective method to ancient human remains from tropical SEA (SOM1). We found a modified version of MYbaits Whole Genome Enrichment to be the best-performing method. We applied this method, in combination with shotgun sequencing approaches where sufficient endogenous DNA allowed it, to samples from Malaysia, Thailand, Philippines, Vietnam, Indonesia and Laos, dating between 0.2 and 8 kya (SOM2). We obtained 25 low-coverage ancient genomes (Table 1), along with mtDNA and nuclear DNA from an additional set of 16 individuals (Table S3), belonging to hunter-gatherers from the Hoabinhian culture, as well as Neolithic, Bronze Age and Iron Age farmers (SOM3). All samples showed damage patterns typical of ancient DNA (*38*) (Table S3).

**Table 1.** Meta-data for ancient samples, including IDs, radiocarbon dates and groups of similar ancestry into which they were placed for reference in the main text. A question mark denotes that a sample is of too low coverage to obtain information about genetic sex or haplogroup.

| Site | ID | Culture | Group | Calibrated date | Genetic Sex | Coverage | mt-haplogroup | Y-haplogroup |
| --- | --- | --- | --- | --- | --- | --- | --- | --- |
| Pha Faen<br>(Laos) | La368 | Hunter-gatherer,<br>Hoabinhian, flexed burial | 1 | 7888 ± 40 | XY | 0.603 | M5 | C |
| Gua Cha (Malaysia) | Ma911 | Phase 1 - Hoabinhian | 1 | 4319 ± 64 | XY | 0.131 | M21b1a | D |
|  | Ma912 | Phase 2 - Neolithic farmer<br>cemetery | 2 | 2447 ± 65 | XY | 1.729 | M13c | O1b1a1a1b1 |
| Ma Dai Dieu<br>(Vietnam) | Vt833 | Neolithic (upper layer) | 2 | 4171 ± 58 | XX | 0.128 | M20 | N/A |
|  | Vt777 | Neolithic (upper layer) | 3 | 2276 ± 62 | XX | 0.147 | F1a1'4 | N/A |
| Tam Hang (Laos) | La898 | Recent intrusion into<br>Hoabinhian | 2 | - | XY | 0.114 | N9a6 | O |
|  | La727 |  | 2 | 2335 ± 14 | XX | 0.942 | N9a6 | N/A |
| Tam Pa Ping (Laos) | La364 | Late Neolithic-Bronze Age | 2 | 2996 ± 47 | XY | 1.451 | F1a1a1 | O |
| Hon Hai Co Tien<br>(Vietnam) | Vt880 | Neolithic - Ha Long<br>Culture | 2 | ~3.5kya | ? | 0.103 | ? | ? |
|  | Vt719 | Intrusive burial | 3* | 229 ± 69 | XX | 0.257 | M7c2 | N/A |
| Nam Tun (Vietnam) | Vt778 | Late Neolithic | 4* | 2652 ± 83 | XY | 0.147 | F1a1a1 | O2a2b1a2a1 |
| Nui Nap (Vietnam) | Vt808 | Dong Son Culture | 3 | 2257 ± 75 | XX | 0.118 | M7b1a1 | N/A |
|  | Vt781 |  | 3 | 2260 ± 64 | XX | 0.139 | F1a | N/A |
|  | Vt779 |  | 3 | 2256 ± 63 | XX | 0.141 | M7c1b2b | N/A |
|  | Vt796 |  | 3 | 2177 ± 116 | XX | 0.110 | F1e3 | N/A |
| Long Long Rak<br>(Thailand) | Th519 | Iron Age | 4 | 1731 ± 51 | XY | 0.161 | B5a1d | N |
|  | Th521 |  | 4 | 1712 ± 59 | XY | 0.422 | F1f | O1b1a1a1b |
|  | Th703 |  | 4 | 1669 ± 36 | XY | 0.163 | B5a1d | NO |
|  | Th531 |  | 3* | 1597 ± 34 | XX | 0.086 | G2b1a | N/A |
|  | Th530 |  | 4 | 1668 ± 36 | XY | 0.196 | G2b1a | IJK |
| Loyang Ujung<br>Karung (Indonesia) | In661 | Late Neolithic-Iron Age,<br>flexed burials | 5 | 1866 ± 26 | XX | 0.105 | F1a1a | N/A |
|  | In662 |  | 5 | 2199 ± 82 | XY | 0.143 | M20 | O1b |
| Nagsabaran<br>(Philippines) | Iron Age | Red-slipped pottery -<br>Austronesian | 6 | 1818 ± 44 | ? | 0.029 | ? | ? |
| Supu Hujung 4 | Ma554 | Historical | 6 | 424 ± 68 | XY | 0.343 | F3b1a+160<br>93 | O |
| Kinabatangan | Ma555 | Historical | 6 | 231 ± 69 | XY | 0.549 | B4b1a2 | O2a2a1a2 |

To address the genetic relationships among the ancient individuals, we performed a principal component analysis (PCA) with our Pan-Asia Panel (see Methods) using *smartpca* (*39*). We projected the ancient samples onto the first two principal components of a PCA constructed solely with present-day samples (*40*) (SOM4). We then used ADMIXTURE (*41*) to find reference latent ancestry components that could best fit our present-day data, and then used *fastNGSadmix (42, 43)* to model the low-coverage ancient samples as mixtures of these reference components (SOM5). Unlike all other ancient samples, the two Hoabinhian samples (which also happen to be the oldest samples in our study) - Pha Faen, Laos (La368 - ^14^C 7,888 ± 40) and Gua Cha, Malaysia (Ma911 - ^14^C 4,319 ± 64) - designated as Group 1, cluster distantly from most East and Southeast Asians in the PCA and position closely to present-day Onge (Figure 1A). Group 1 individuals also contain a mixture of several different ancestral components in the *fastNGSadmix* plot, including components shared with Onge, the Pahari and Spiti from India, Papuans and Jehai (a Malaysian ‘Negrito’ group), which are markedly different from the other SEA ancient samples. This possibly results from our modeling of ancient populations as a mixture of components inferred in present-day populations, via *fastNGSadmix* (*44*), and from the fact the ancient samples are likely poorly represented by a single present-day group. The rest of the ancient samples are defined primarily by East and Southeast Asian components that are maximised in present-day Austroasiatic (Mlabri and Htin), Austronesian (Ami) and Hmong (indigenous to the mountainous regions of China, Vietnam, Laos and Thailand) populations, along with a broad East Asian component.

**Figure 1.**
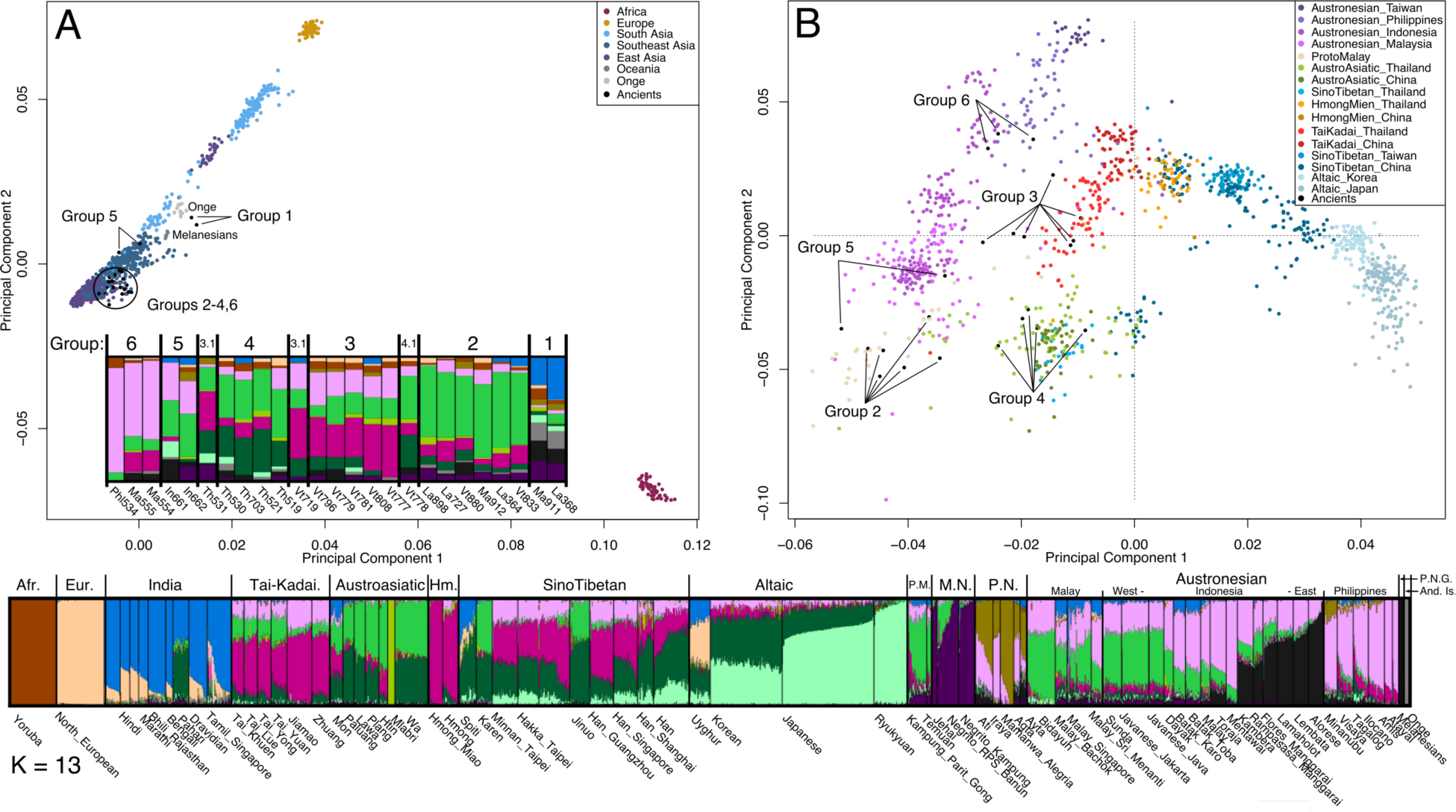
A) First two components of PCA of world-wide populations, including ancient SEA individual projections, computed using the Pan-Asia panel data. B) First two components of PCA of present-day and ancient individuals from mainland SEA, excluding Onge and the ancient Hoabinhians (Group 1), highlighting the differences in ancestry affinities among individuals from Groups 2-5. Inset of panel A: *fastNGSadmix* plot for ancient samples, classified into groups of similar ancestry. Lower panel: *fastNGSadmix* plot at K=13, for all present-day samples, excluding SGDP genomes (see SOM5). We refer to the following present-day language speaking groups in relation to our ancient samples - Austroasiatic (Mlabri and Htin - bright green), Austronesian (Ami - pink) and Hmong (indigenous to the mountainous regions of China, Vietnam, Laos and Thailand - dark pink), along with a broad East Asian component (dark green).(Hm=Hmong Mien, P.M.=Proto Malay, MN = Malaysian ‘Negrito’, PN = Philippines ‘Negrito’, P.N.G. = Papua New Guinea, And. Is. = Andaman Islands)

We used outgroup f_3_ statistics (f_3_(Mbuti;X,Ancient samples)) to determine which populations have the highest levels of shared drift with each of the ancient individuals (SOM6). Group 1 shares the most drift with certain ancient mainland samples (Figure S12, Table S4). Again, we see that the closest present-day populations to Group 1 are from the Andaman Islands (Onge) and then Kensiu (a Malaysian ‘Negrito’ group), Ami and Jehai, followed by a mix of East and Southeast Asian populations.

We used D-statistics of the form D(Papuan,Tianyuan,X,Mbuti), where X is a test population, to explore the relatedness of ancient and present-day Southeast Asians to two highly differentiated groups: Papuans and an ancient northern East Asian individual (Tianyuan - a 40 kya-old sample from Northeastern China (*45*)). The values of this D-statistic are consistent with present-day and ancient SEA mainland samples being more closely related to Tianyuan than to Papuans (SOM7). This applies to present-day northern EA populations, and - more weakly - to most populations of ancient and present-day SEA. However, this D-statistic is not significantly different from 0 in present-day Jehai, Onge, Jarawa and Group 1 - the ancient Hoabinhians (Figure 2B, Tables S12, SOM7). While the Onge’s relationship with Papuans and Tianyuan is unclear, D-statistics of the form D(Onge,Tianyuan,X,Mbuti), where X is a test population, show that Jarawa, Jehai and the ancient Group 1 share more ancestry with Onge than with Tianyuan (Figure 2C, SOM7). Like the Onge, both Group 1 samples carry mtDNA haplogroups from the M lineage (Table S3), thought to represent the coastal migration to Australasia (*12, 13, 28, 46*).

**Figure 2.**
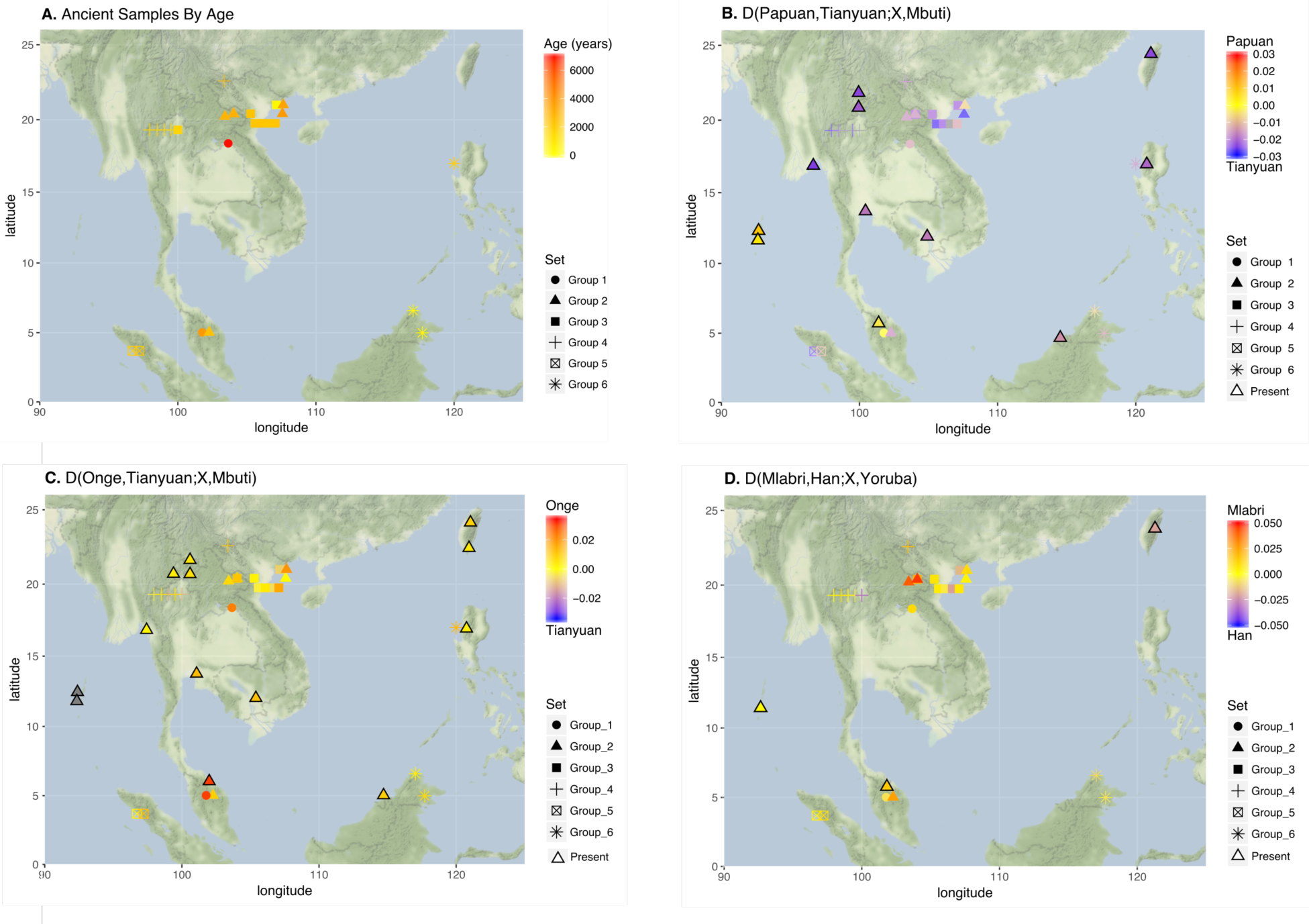
A) Estimated mean sample ages for ancient individuals. B) D-statistics testing for differential affinity between present-day Papuans and Tianyuan (2240k Panel). C) D-statistics testing for differential affinity between present-day Onge and Tianyuan (2240k Panel). D) D-statistics testing for differential affinity between present-day Mlabri and present-day Han Chinese (Pan-Asia Panel).

To assess the diversity among the remaining ancient individuals, we computed a new PCA including only EA and SEA populations that did not have considerable Papuan or Onge-like ancestry in the *fastNGSadmix* analysis (Figure S11), as it was done in the Pan-Asian SNP-capture study (*30*). We observe that the remaining ancient samples form five slightly differentiated clusters within the EA and SEA populations (Groups 2-6, Figure 1B), in broad concordance with the *fastNGSadmix* (at K=13, Figure 1) and f_3_ results (Figure S12-S19; SOM4). We thus decided to organize these samples into five more groups to facilitate further analyses (Groups 2-6, Table 1), although we note that genetic differentiation among them seems to be highly clinal. Samples Vt719, Th531 and Vt778 were either geographic or temporal outliers to their groups and were therefore analyzed separately in groups denoted by a “.1”: Group 3.1 (Th531, Vt719) and Group 4.1 (Vt778).

Group 2 samples from Vietnam, Laos, and the Malay Peninsula are the oldest samples after Group 1, and range in age from 4.2 to 2.2 kya. At K = 6 (SOM5), Group 2 individuals, the present-day Mlabri and a single Htin individual are the only MSEA samples in the *fastNGSadmix* analysis to lack the broad EA component (dark green) maximised in northern EA, with the exception of the Malaysian ‘Negritos’ and ‘Proto-Malays’ (Temuan). At K = 7, a bright green component is maximised in these populations, and this component is also found in present-day Indonesian samples west of Wallace’s Line. The two ancient Indonesian samples (Group 5; 2.2 to 1.9 kya) represent a mix of Austronesian-and Austroasiatic-like ancestry, similar to present-day western Indonesians. Indeed, after Mlabri and Htin, the closest populations to Group 2 based on outgroup-f3 statistics are the western Indonesian samples (from Bali and Java) reported to have the highest amounts of ancestry from mainland SEA (*47*) (Figure S13).

These lines of evidence suggest Group 2 are possible descendants of an “Austroasiatic” migration that expanded southward across MSEA and into island SEA (ISEA) by 4 kya (*27, 47*–*49*). We also observe a gradient in “Austronesian-like” vs. “Austroasiatic-like” ancestry in the PCA (Figure 1B): while PC1 separates populations found in SEA and those found in northern EA, PC2 distinguishes population based on their amounts of Austronesian-like ancestry (pink component in Figure 1 - lower panel) versus Austroasiatic-like ancestry (bright green component in Figure 1 - lower panel).

Group 6 samples are recent (between 1.8 and 0.2 kya) and come from Malaysia and the Philippines. They fall within the variation of present-day populations with high Austronesian ancestry in these areas. Group 6 also contains the individual (Ma554) with the highest amounts of Denisovan ancestry relative to the other ancient samples, although variation in archaic ancestry is not very strong across MSEA (SOM10).

The remaining mainland samples (Groups 3 and 4) are dated to be from 2.6 to 0.2 kya. They appear similar to present-day MSEA populations and fall into two groups. Group 3 is largely composed of ancient samples from Vietnam but also includes one sample from Thailand (Th531); these samples cluster in the PCA with the Dai from China, Tai-Kadai speakers from Thailand and Austroasiatic speakers from Vietnam, including the Kinh (Figures S9). In contrast, Group 4 largely contains ancient samples from Long Long Rak, Thailand, but also includes the inland-most sample from Vietnam (Vt778). These samples fall within the variation of present-day Austroasiatic and Sino-Tibetan speakers from Thailand and China, supporting the hypothesis that the Long Long Rak population originated in South China, and subsequently expanded southward during the Dongson period (*50*). At Long Long Rak, three individuals (Th387,Th530 and Th531) dated to approximately 1.6 kya were found in the same chamber. Interestingly, all three individuals share the same mtDNA haplogroup (G2b1a), but the nuclear ancestries for the two samples which yielded genome-wide data are quite different: Th531 clusters best with Group 3, while Th530 with Group 4. These results suggests that individuals with ancestry from distant regions likely cohabited at this locality.

To determine if any of the ancient samples had affinities to particular populations outside SEA, we computed D-statistics of the form D(Group A, Group B, Not-SEA, Yoruba/Mbuti) to compare each of the ancient groups. Group 2 has a significant affinity to the Indian populations of Khonda Dora (Z = 6.3), relative to Group 3 (D(Group2,Group3,Khonda Dora,Yoruba)), in agreement with previous reports of East Asian ancestry in tribal Indian Groups (*51, 52*). We also investigated the affinity between certain Australasian populations and particular Native American groups, like the Surui (*45, 53, 54*). When computing D(Mixe, Surui, X,Yoruba), we find that the Group 1 samples had some suggestive but non-significant affinity to Surui relative to Mixe (Z = -2.18 when X = Ma911, Z = -2.48 when X = La368; Table S19), although the signal is not as robust as observed for Tianyuan (Z = -3.53), Khonda Dora (Z = -3.04) and Papuans (Z = -3.02), among others (*53, 54*). We note, however, that there are much fewer SNPs to compute this statistic on Group 1 samples than on the other populations (La368: 191,797; Ma911: 47,816; Tianyuan: 295,628, Papuan: 471,703, Khonda Dora: 496,097), thus we may be underpowered to detect this signal.

We used *TreeMix* (*55*) to explore admixture graphs that could potentially fit our data. The ancient Group 1 (Onge-like) individuals are best modelled as a sister group to present-day Onge (Figures 3A, S21-S23). For the highest-coverage Group 1 sample, allowing for one migration, TreeMix fits Papuans as receiving admixture from Denisovans, while the second migration shows East Asian populations as resulting from admixture between Tianyuan and Onge.

**Figure 3.**
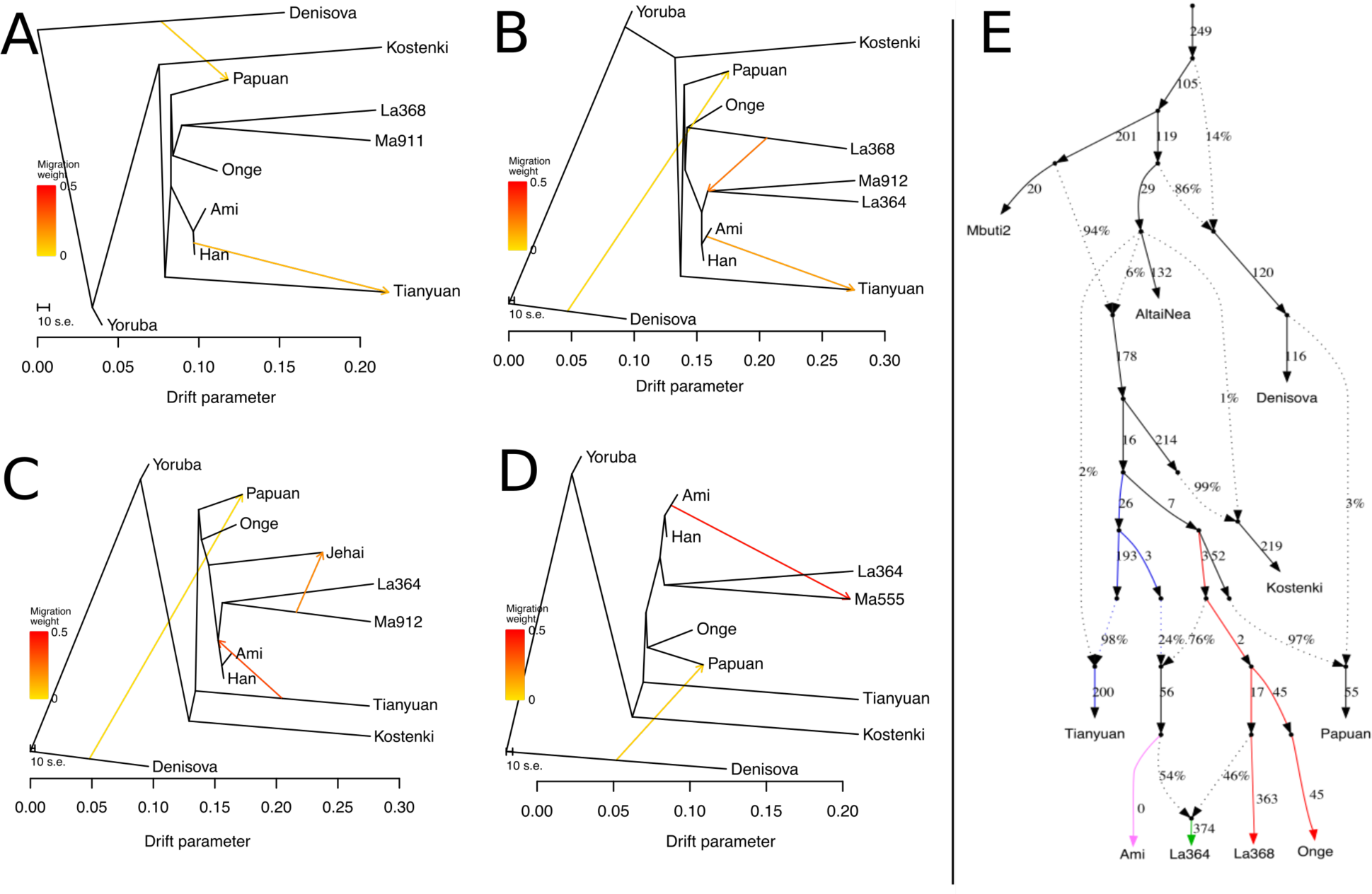
We used TreeMix to build admixture graphs combining present-day populations and select ancient samples with high SNP coverage. A) A graph including both Group 1 samples (Ma911 and La368) shows they can be fitted as sister groups with close affinities to present-day Onge. B) A graph including the highest coverage Group 1 (La368) individual and Group 2 (La368, Ma912) samples is best fit with Group 2 receiving ancestry from both Group 1 and the East Asian branch. C) A graph including the Group 2 individual and Jehai, showing admixture between Jehai and Ma912. D) TreeMix models Ma555 (Group 6) as receiving ancestry from both a branch leading to La364 (Group 2) and present-day Igorot. E) We used a graph framework inferred by Lipson et al. (*61*) and attempted to fit different ancient and present-day SEA individuals within that framework in *qpGraph*. Note that we model present-day East Asians (here represented by Ami) as a mixture of an Onge-like population and a population related to the Tianyuan ancient individual. La368 is best fitted as a sister group to Onge, while La364 is best fitted as a mixture of the ancient Onge-like population (represented by La368) and an East Asian population (represented by Ami) (worst-fitting Z = 3.667).

We also performed a more supervised form of admixture graph modeling using *qpGraph (36)* (SOM9). We began with a skeletal framework containing chimpanzee (PanTro2, EPO alignment from Ensemb 71 (*56, 57*)), Denisova (*58*), Altai Neanderthal (*59*), Kostenki-14 (*60*), Mbuti, Onge, Ami and Papuan, fitting a graph based on results from Lipson and Reich (*61*) and well-supported D-statistics (SOM7). When not including Tianyuan, we find that the Onge-Papuan-Ami split is hard to resolve (Figure S33), in agreement with Lipson and Reich (*61*). However, when including Tianyuan (Figure S34), we find that a best fit occurs when Ami (or Han) are modelled as an admixed group, with ancestry from a population related to Tianyuan and a population related to Onge (worst-fitting Z = -3.564). In support of this graph assignment, we find that D(Ami, Onge, Tianyuan, Mbuti) = 0.0239 (Z = 5.148), while Papuan and Onge are a clade with respect to Tianyuan: D(Papuan, Onge, Tianyuan, Mbuti) = -0.0047 (Z = -0.886). We then added either La368 or Ma911 to the graph. In agreement with the *TreeMix* results, we find that La368 and Ma911 are each best modeled as a sister group to Onge (Figures S35 and S36, worst-fitting Z = 3.372 and 3.803, respectively).

We then used *qpWave/qpAdm (62, 63)* to determine if La368 and Ma911 can be modelled as a linear combination of ancestries from Papuans, Onge and/or Tianyuan without the need to invoke partial ancestry from a population that may have split from them before these populations split from each other. As outgroup populations, we used Yoruba (*64*), Ust-Ishim (*65*), Kostenki-14 (*60*), Mal’ta (*66*), Afontova Gora 3, Vestonice 16, El Mirón and Villabruna (*67*). All best 3-way and 2-way combinations for La368 are not feasible (have negative admixture weights). There are two 1-way possibilities (La368 as a sister group to either Onge or Papuans) that are feasible and are good fits (P = 0.37 and P = 0.27, respectively), and this is somewhat expected as Onge and Papuans are sister clades to each other - barring Denisovan introgression into Papuans. When performing the same analysis on Ma911 as the target population, we find that all the best 3-way and 2-way combinations are also infeasible and the only good 1-way fit is with Onge (P = 0.49). Modelling Ami as a linear combination of the same three source populations results in any of the 3-, 2- or 1-way fits being feasible and good fits, but the best fit is found in the 2-way combination of Tianyuan and Onge (P = 0.98).

When modeling the Mlabri-like Group 2 in *TreeMix*, we see that the two samples with the highest coverage in this group (La364 and Ma912) form a clade, resulting from an admixture event between the ancestral populations of present-day East Asians (Han/Ami) and the ancestors of La368 (Figures 3B, S24-27). Despite the low SNP overlap (∼20,000 SNPs) when including the Group 1 and 2 samples from Laos and Malaysia, (La368, Ma911, La364, Ma912), at 3 migrations, *TreeMix* residuals suggest that the Onge-like ancestry in Malaysia and Laos is a result of local admixture (Figure S27, SOM8). Additional data and higher coverage samples from these regions are needed to better support a ‘local admixture’ model: including all four low-depth genomes in the same admixture graph results in only 17,286 overlapping SNPs (including transitions), which makes inference difficult. The Jehai are best fitted as an admixed population between Group 2 (Ma912) and the branch leading to present-day Onge and La368 (Figure 3C, S28). ISEA ancient samples from Indonesia (Group 5) and Borneo (Group 6) are best modelled as an admixed population carrying the signature of Group 2 (Figure 3D, Figures S29-33), supporting a previously reported mainland component in ISEA complementary to the well-documented Austronesian expansion (*47*). For the ISEA samples (Group 5 - In662 and Group 6 - Ma554), when a more basal migration event occurs, it originates from the Papuan branch, rather than the Onge branch as seen in MSEA.

Consistent with the *TreeMix* results, La364 in *qpGraph* is best modeled as a mixture of a population ancestral to Ami and the Group 1 / Onge-like population (Figure 3E, worst-fitting Z = 3.667). Additionally, we find the best model for present-day Dai populations is a mixture of Group 2 individuals and an additional pulse of admixture from East Asians (Figure S37, worst-fitting Z = 3.66).

This is the first study to reconstruct the population history of SEA using ancient DNA. We find that the genetic diversity found in present day SEA populations derives from at least four prehistoric population movements by the Hoabinhians, an “Austroasiatic-like” population, the Austronesians and, finally, additional EA populations into MSEA. We further show that the ancient mainland Hoabinhians (Group 1) shared ancestry with present-day Onge of the Andaman Islands and the Jehai of peninsular Malaysia. These results, together with the absence of significant Denisovan ancestry in these populations, suggest that the Denisovan admixture observed in Papuans occurred after their ancestors split from the ancestors of the Onge, Jehai and the ancient Hoabinhians. This is also consistent with the presence of substantial Denisovan admixture in the Mamanwa from the Philippines, which are best modeled as resulting from an admixture between Austronesians and Papuans, not Onge (*61*).

Consistent with the Two Layer model, we observe a dramatic change in ancestry by 4 kya (Group 2) which coincides with the introduction of farming, and thus supports models that posit a significant demographic expansion from EA into SEA during the Neolithic transition. Group 2 are the oldest samples with distinctive EA ancestry that we find. The most closely related present-day populations to Group 2 are the Mlabri and Htin - the Austroasiatic hill tribes of Thailand - which is in agreement with hypotheses of an early Austroasiatic farmer expansion into the region. They also share ancestry with the Temuan and Jehai of Peninsular Malaysia and populations of Western Indonesia, supporting an Austroasiatic (“Western Route”) expansion into ISEA (post-Hoabinhian, pre-Austronesian), as previously proposed based on linguistic and archaeological grounds (*27, 49, 68*). Furthermore, a recent study also identified populations of Bali and Java as the groups in ISEA with the highest frequency of mainland SEA ancestry (*47*), also reflected in the large amounts of shared drift between Group 2 and the Javanese that we observe (Figure S13). The extent and nature of this Austroasiatic expansion into western Indonesia prior to the Austronesian expansion could be resolved by sequencing ancient genomes from ISEA prior to the Austronesian expansion.

By around 2 kya, all ancient mainland samples carry additional EA ancestry components that are absent in Group 2. Within the variation of these recent samples, we find two clusters of ancestry, possibly representing independent EA migrations into mainland SEA. Group 3 has affinities to the Hmong, the Dai from China, the Thai from Thailand and the Kinh from Vietnam, while Group 4 individuals - found only in inland regions - have affinities to Austroasiatic Thai and Chinese speakers. Finally, we also find evidence for the arrival of Austronesian ancestry into the Philippines by 1.8 kya (Group 6) and into Indonesia by 2.1 kya (Group 5). By 2 kya, the population structure in MSEA was very similar to that among present-day individuals. Despite observing a clear change in genetic structure coinciding with the transition from the Hoabinhian hunter-gatherers to Neolithic farmers, we also see a degree of local continuity at all sites at different points in time, suggesting that incoming waves of migration did not completely replace the previous occupants in each area (Figure 4).

**Figure 4.**
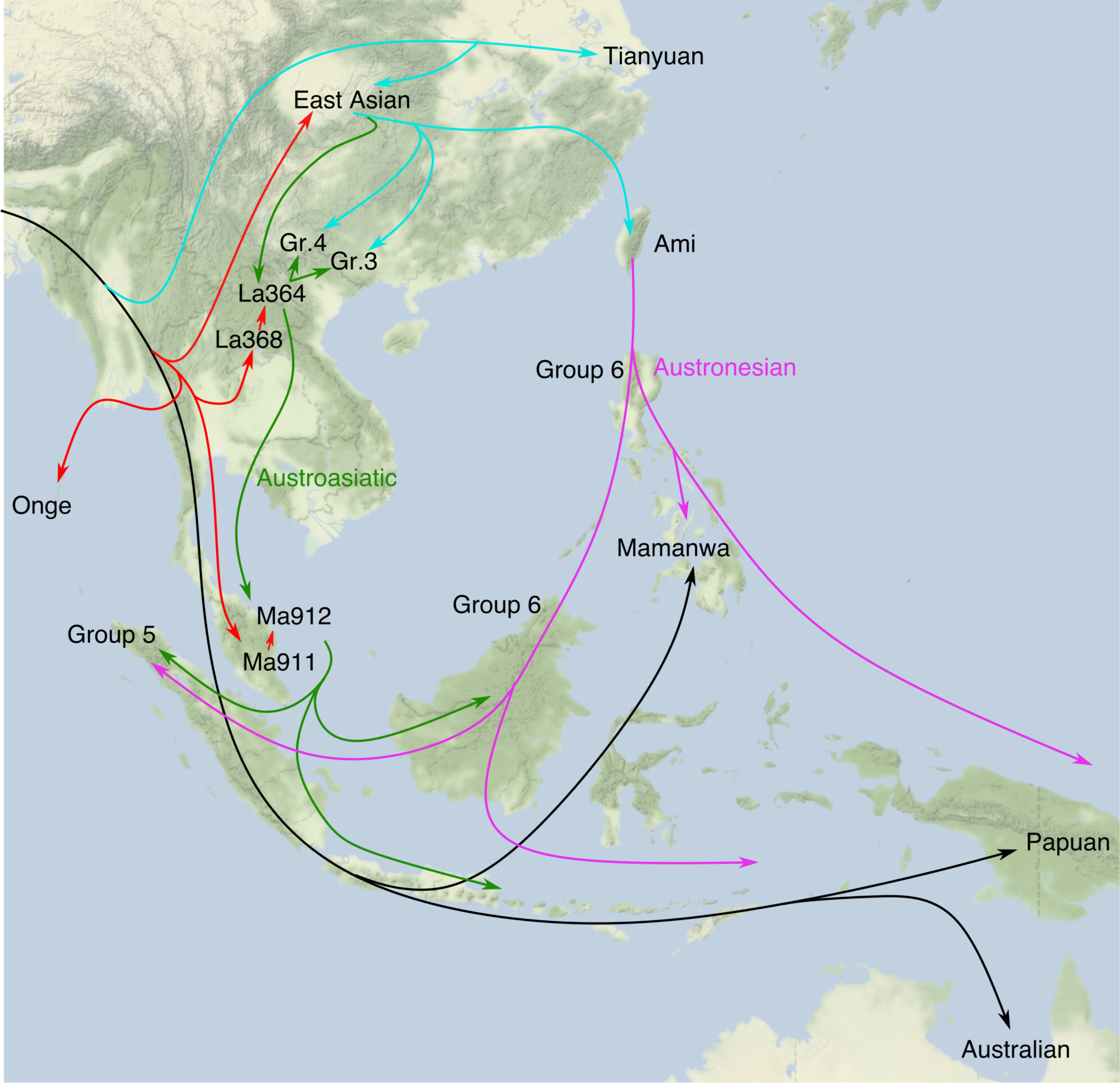
A model for plausible migration routes into Southeast Asia, based on the ancestry patterns observed in the ancient genomes.

This study demonstrates that whole-genome capture is an efficient supplementary approach for retrieving whole genomes from the fossil skeletal and dental remains found in the tropics. As target enrichment inevitably results in subsampling library fragments, it is most useful for (combined) libraries with high underlying complexity. We found a median 7.5 fold enrichment, reducing the sequencing costs proportionally. By enriching the human DNA content, we were able to acquire whole-genome data from selected samples in which the low proportion of endogenous DNA would have been previously prohibitive. The whole genome approach which we have employed here combines shotgun sequencing and capture in order to maximise the potential of ancient samples.

The clear genetic distinction between the Onge-like Hoabinhian and EA Neolithic demonstrated by this study provides an overwhelming support for the Two Layer model and indicates that in SEA, like in Europe, the onset of agriculture was accompanied by a demographic transition. However, on a more local level, our results point toward admixture events in northern Laos and Peninsula Malaysia between the two dispersal layers. We also show that the Hoabinhians of the first dispersal contributed a degree of ancestry to the incoming EA populations, which may have also resulted in the passing on of some phenotypic characteristics detected by proponents of the Continuity model. Finally, our results reveal that the appearance of these Austroasiatic farmers at around 4 kya was followed by multiple migrations of distinct EA ancestry. These subsequent migrations made significant contributions to the diversity of human populations in present-day SEA.

## Methods

### Samples

We screened ancient samples from across SEA. We prioritized petrous bone, because of its favorable DNA preservation (*69*). Most of the samples were processed at the Centre for GeoGenetics, University of Copenhagen, with a few at Griffith University (Long Long Rak, Thailand; Th387, Th391, Th392, Th389, Th126, Th127, Th238, Th248), in dedicated clean laboratories following strict ancient DNA guidelines (*69*–*71*). Material was sampled as described in Hansen et al. (*69*). To minimize risk of contamination from handling, we performed a pre-digestion step (*72*). DNA extraction was done as in (*73*) followed by dual-indexed libraries building and amplification (*74*). Adapter-dimers were removed where necessary, using AMPure beads.

### Sequencing, Mapping and Genotyping

Sequencing was performed on Illumina Hiseq2500 (ver. 4) or Hiseq4000 instruments (81bp single-read) using bcl2fastq de-multiplexing. Adapters were trimmed using *AdapterRemoval* v2.2.2 (*75*), and mapped to the human reference genome (hg19, build 37) using BWA (*76, 77*) (SOM3).

To minimize batch effects, we obtained genotypes for all individuals from a combination of published BAM files from previous studies and BAM files produced in this study. We genotyped genomes that had low coverage or were obtained from targeted capture by selecting the majority allele for the genomic position, looking only at reads with mapping quality ≥ 30 and base quality ≥ 30. If both alleles were present at a site with equal coverage, a random allele was selected. High-coverage genomes were genotyped as in Sikora et al. (*78*). All analyses were restricted to regions that were within the 1000 Genomes Phase 3 (*79*) strict accessibility mask (ftp://ftp.1000genomes.ebi.ac.uk/vol1/ftp/release/20130502/supporting/accessible_genome_ma sks/20141020.strict_mask.whole_genome.bed), and outside repeat regions (UCSC genome browser simpleRepeat table).

### Damage/Contamination

Using *mapDamage* v2 *(80)*, we verified that all ancient samples displayed signatures of cytosine deamination and short fragment lengths, both typical of ancient DNA. For all samples in Table 1, contamination estimates obtained using *contamMix* (*65*) were minimal (MAP probability of authentic ancient DNA material = 0.94-0.99%, depending on the sample), while some of the very low coverage samples that were not sequenced to genome-wide depth had elevated contamination rates (Table S3). When practical, extracts were USER-treated (*81*) for deep sequencing after damage patterns were identified from screening results.

### Reference Panels

We assembled two panels for different types of analyses. Initial analyses were undertaken using the HUGO Pan-Asian SNP Database (*30*) (1,744 individuals; 50,796 SNPs) and Onge from the Simons Genome Diversity Panel (SGDP) (*64*), resulting in a panel maximising populations, at the expense of a lower SNP number, with 50,136 overlapping SNPs (hereafter the “Pan-Asia panel”). We assembled a second panel using whole genomes from the SGDP, limiting to the 2,240k capture SNPs from Yang et al. (*45*). We used the first panel for ADMIXTURE / *fastNGSadmix* analyses and PCA, as well as f3 statistics. We used the second panel for more parameter-rich modelling.

### Principal Component Analysis

We performed a principal component analysis using *smartpca* v1600 implemented in the *Eigensoft* package (*82*) on the SNP covariance matrix of genomes from the Pan-Asia panel. We then projected all our ancient SEA samples onto the first two principal components (*40*) (SOM4).

## ADMIXTURE

We ran ADMIXTURE v1.3.0 (*41*) from K = 1 to K = 13 on the Pan-Asia Panel and the SGDP panel, after LD-pruning in PLINK (*83*), yielding 35,042 SNPs for analysis. To get standard errors for parameter estimates, we obtained 200 bootstrap replicates in each run. We then modeled low-coverage ancient populations based on the reference components inferred by ADMIXTURE, using *fastNGSadmix* (*43*). To visualise the admixture plots we used *pong* (*84*). Throughout this study, we generally refer to the colors corresponding to the ancestry components assuming K=13, unless otherwise stated.

### *f* and *D* statistics

We computed *f* and *D* statistics to measure the amount of shared drift between two populations, and to test gene-flow and treeness hypotheses, as detailed in Patterson et al. (*36*). For both analyses, we estimated standard errors through a weighted block jackknife procedure over 5Mb-blocks. For *D* statistics, we restricted the analysis to transversion polymorphisms in order to minimize potential bias introduced by differential error rates in ancient samples (mostly a consequence of *post-mortem* ancient DNA damage and low depth).

### TreeMix

We performed unsupervised admixture graph fitting of our ancient and present-day samples using *TreeMix* v1.13 (*55*) on the data. We used high coverage genomes from Denisovan, Mbuti, Kostenki, Papuan, Onge, Papuan, Tianyuan, Han and Ami as a base set of populations, and rooted the graph using Denisova as the outgroup. For each test, we only considered sites where all analysed populations had at least one individual with non-missing data and grouped SNPs in 5Mb blocks (-k parameter) to account for linkage disequilibrium. We observed that including transitions for ancient samples caused biases in inference and so removed transitions in all analyses. We show all graphs fitted in SOM8.

### qpGraph

We ran *qpGraph* v6100 from the *Admixtools* package (*36*), following parameter settings as in Lipson and Reich (*61*). We used fitted graphs with chimpanzee at the root, set “outpop” to be “NULL” to prevent use from specifying a particular outgroup population in which SNPs must be polymorphic, used a block size of 0.05 Morgans for the jackknife procedure, and used the full matrix form of the objective function, with “diag” set to 0.0001. Finally, we set a Z-score = 3 as the cutoff to label a statistic as an outlier. For further details, see SOM9.

## Author Contributions

EW initiated the study. EW, DML, LV, LO, HM and FD designed the study. EW and DML supervised the overall project, while LV, FD, FR, VS, TS, MMS, RS, HMN, CH, KW, EPE, JCG, RK, HB and CP supervised specific aspects of the project. HM, LV, FD, UGW, CD, VS, TS, MMS, RS, SK, PL, HMN, HCH, TMT, THN, SS, KW, AMB, PD, JLP, LS, EPE, NAT, JCG, RK, HB and CP excavated, curated, sampled and/or described samples. HM, LV, ASO, SW, PBD, SR and TH produced data for analysis. HM, FR, LV, JVMM, CD, SW, AM, LO and MS analysed or assisted in the analysis of data. HM, FR, LV, FD, AM, LO, MS, CH, DML and EW interpreted the results. HM, FR, LV, FD, MML, RAF, CH, DML, EW wrote the manuscript with considerable input from JVMM, CD, SW, AP, VS, TS, MMS, RS, HMN, HCH, THN, KW, TH and MS. All authors discussed the results and contributed to the final manuscript.

## Acknowledgements

We thank the National High-throughput DNA Sequencing Centre (Copenhagen Denmark) for expert advice and sequencing of samples, the Duckworth Laboratory, University of Cambridge, for permission to sample material in their care, and Kristian Gregersen for making casts of teeth before sampling. This work was supported by the Lundbeck Foundation, the Danish National Research Foundation, and the KU2016 program. HM is supported by the George Murray Scholarship (University of Adelaide). RS thanks the Thailand Research Fund (TRF) for their support (Grants RTA6080001 and RDG55H0006). MML is supported by the ERC award 295907. DML was supported by ARC Grants LP120200144, LP150100583 and DP170101313. EW thanks St. John’s College, University of Cambridge, for providing an inspiring environment for scientific thought.

## List of Supplementary Materials

SOM1. Assessment of target enrichment methods

SOM2. Archaeological Overview

SOM3. Mapping

SOM4. Principal Component Analysis

SOM5. ADMIXTURE fitting

SOM6. f3 Statistics

SOM7. D-statistics

SOM8. TreeMix fitting

SOM9. qpGraph fitting

SOM10. Measurements of archaic ancestry

Table S1 - S19

Fig S1 - S41

Supplementary References: 89-115

